# Influenza Virus-Induced Autophagy Is Not Always Associated with MUC1 Expressions

**DOI:** 10.1101/2022.05.25.492935

**Authors:** Hasan Selcuk Ozkan, Mustafa Alper Ozarslan, Gokhan Vatansever, Candan Cicek

## Abstract

Influenza virus-induced autophagy frequently accompanies apoptosis and results in cell death in cells infected with the virus. Autophagy has been well-known to be modulated by the mTOR/PI3K/Akt pathway, which plays an important role in response to the presence of energy sources and external stimulants. This pathway can also be modulated by MUC1, which has extracellular and intracellular components, and playing an important role in metastasis and chemotherapeutic resistance. In this study, the aim is to observe the changes in MUC1 expressions, which is known to have sialic acid residues within MUC1, therefore serving as a receptor for influenza viruses, and consequent changes in autophagy markers such as mTOR and LC3b, after inoculation of cancer cells with influenza virus.

Fluorescence was detected for LC3b, mTOR and MUC1 in all influenza-positive cell lines (MCF-7, He-La, A-549) except for RD in immunofluorescence studies. In influenza-negative MCF-7, HeLa, RD and A-549 cells no fluorescence was detected. In the supernatant of all influenza-positive cells, except for RD cells, positive results were obtained for *MUC1, MAP1LC3B* and *MTOR* genes (Ct 20,01-32,04) in RT-qPCR, respectively encoding for MUC1, LC3b, and mTOR. In RT-qPCR, in cell lines without influenza inoculation, only in the A-549 cells, the gene expressions were found to be negative. In other cell lines, positive results were obtained (Ct 21,98-23,78). In RD cells, which have not been inoculated with the virus, all pathways were positive.

In cells with adenocarcinoma structure, the alterations and presence of autophagy pathways and MUC1 expressions activated via influenza viruses were confirmed. However, this has not been proved using nucleic acid assays. The reason why might be the detection of only gene expressions and mRNA presence via the tests used in the study. In future studies, expressions of these genes in protein level should be detected using more advanced tests.

## 1. Introduction

Influenza virus-induced autophagy is a prerequisite for the apoptosis of influenza infected cells and has some specific characteristics when compared to the traditional autophagy, as it is often accompanied by increased gene expressions of mammalian target of rapamycin (mTOR), during the late-stage replication of the virus^1,2^. Mucins, on the other hand, are one of the most important components of the innate immunity as they provide the physical barrier of the epithelia against the intraluminal insults. However, one of these structures, Mucin 1 (MUC1) doesn’t only serve to this function as it consists of both intracellular and extracellular components, therefore leading to different distribution patterns in cancer cells due to the loss of apical-basal polarity, yielding to prognostic and therapeutic consequences^3,4^, but also serving as a potential target of the influenza virus, as it has sialic acid residues in its extracellular component, which are known to have different glycosylation patterns in cancer cells^5^.

In this study, the aim was to observe the changes in *MUC1* gene expressions in four influenza virus-inoculated cancer cell lines including A-549, MCF-7, HeLa and RD, and consequent changes in survival and autophagy associated genes encoding mTOR and LC3b, as it’s well-established that influenza entry requires the presence of sialic acid residues on the outer layer of the cell membrane^6^.

## 2. Materials & Methods

### 2.1. Cell Culture

A-549 lung adenocarcinoma, MCF-7 breast adenocarcinoma (ATCC, Manassas, VA, USA), HeLa cervix adenocarcinoma, RD rhabdomyosarcoma, and MDCK Madin-Darby canine kidney cells (DSMZ, Braunschweig, Germany) were cultured in modified Eagle’s serum (MEM) supplemented with 10% inactivated fetal calf serum (FCS), 2% L-Glutamine, 1% Penicilline/Streptomycin, 0.1% Gentamicin and 0.4% Amphotericin B and 1M 4-(2-hydroxyethyl)-1-piperazineethanesulfonic acid (HEPES) (Cegrogen Biotech, Ebsdorfergrund, Germany). Cell lines were maintained at 37°C in a 95% humidified incubator with 5% CO_2_. The experiments were performed between passages four to seven. Table 1 summarizes the groups included in the study and the applications to these cells.

**Table 1.**
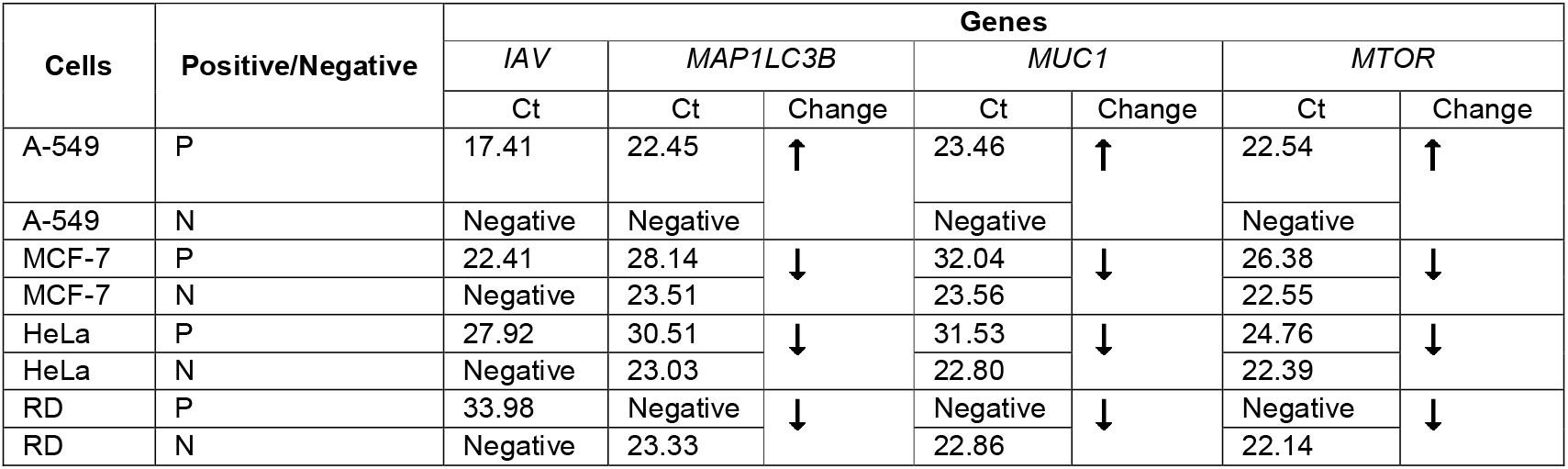
Ct values obtained in qRT-PCR and changes among groups

### 2.2. Virus inoculation

Each cell line (A-549, MCF-7, HeLa, RD) was passaged to 8 shell-vials, after the attachment of cells on the vials, four of the shell-vials were inoculated with the influenza-positive sample, with the other four being left without inoculation as influenza-negative controls. For each reaction two MDCK cell lines passaged, with one being inoculated with influenza, and the other left without, to serve as positive-controls for our virus sample.

To ensure the inoculation of the cells, the supernatant over the vials were removed and the cell-layer was washed twice with MEM supplied with 1% Penicilline/Streptomicin, 0.1% Gentamicin and 0.4% Amphotericin B (Cegrogen Biotech, Ebsdorfergrund, Germany). After adding 0.2 ml of virus suspension to each sample, the plate was centrifuged in 2700g for 30 mins. The supernatant was removed after an hour of incubation at 37^0^C in a 95% humidified incubator with 5% CO_2_. 1 ml of an isolation medium consisting of MEM supplied with 2% L-Glutamine, 1% HEPES, 0.2% trypsin TPCK, 1% Penicilline/Streptomycin, 0.1% Gentamicin and 0.4% Amphotericin B (Cegrogen Biotech, Ebsdorfergrund, Germany). All cells were incubated at 37^0^C in a 95% humidified incubator with 5% CO_2_ for 48 hours.

### 2.3. qRT-PCR

#### 2.3.1. Virus detection from supernatant

Supernatants of all the cells were collected after the end of 48h incubation. Following total RNA extraction from supernatants by RNeasy Mini Kit (Qiagen, Dusseldorf, Germany) and cDNA synthesis by RT^2^ First Strand Kit (Qiagen, Dusseldorf, Germany), presence of influenza nucleic acid was detected using quantitative real-time polymerase chain reaction (qRT-PCR) by a respiratory pathogen detection kit (FTD Diagnostics, Qiagen, Dusseldorf, Germany). The reaction was run on and screened via Rotor-Gene Q (Qiagen, Dusseldorf, Germany).

#### 2.3.2. Gene Expression Detection

After removal of attached cells from the plate using 1 ml MEM supplied with 2μg/ml trypsin TPCK, the buffers are added and centrifuged, then the RNA extraction performed using the RNeasy spin column (Qiagen, Dusseldorf, Germany), as instructed by the producer. The reaction was run on and screened via Rotor-Gene Q (Qiagen, Dusseldorf, Germany), with using the primers for *MUC1, MAP1LC3b* and *MTOR* (Bio-Rad Laboratories, California, USA). All Ct values above ≥30 was considered as weak positive.

### 2.4. Immunofluorescence staining

After fixation of cells with 4% paraformaldehyde, all cultured cells (A-549, MCF-7, HeLa, RD) were stained after the achievement of permeabilization using 0.1% Triton X-100, followed by application of anti-rabbit primary antibodies targeting MUC1, LC3b, mTOR (Abcam, Cambridge, United Kingdom) and an FITC conjugated antibody targeting influenza A and B strains (Light Diagnostics™ MERCK, Darmstadt, Germany) in conformity with the proposed protocols by the producers. All the primary antibodies were coupled with the secondary goat anti-Rabbit antibody Alexa Fluor 488 (Abcam, Cambridge, United Kingdom). Images were obtained via fluorescence microscopy (Nikon, Japan). In all cells observed, one or more fluorescent stained cells were considered as expression of the respective structure.

## 3. Results

In all adenocancer cell lines (A-549, MCF-7, and HeLa) inoculated with influenza virus, antibody staining was found to be positive for MUC1, mTOR and LC3b. However, in all cells not inoculated with the influenza samples and in RD cell line, which is of mesenchymal origin, immunofluorescence was not detected.

In each repeat of the experiments, a total of twice, MDCK cells were also inoculated using the same influenza virus sample, as these cells are known to be a good host to produce influenza viruses, as shown by the many studies conducted in the literature, to ensure the presence of infective viruses inside the cells, whose supernatant was previously positive for influenza nucleic acid. All immunofluorescence staining can be seen in Figures 1 and 2, respectively for influenza-positive and negative cells.

**Figure 1.**
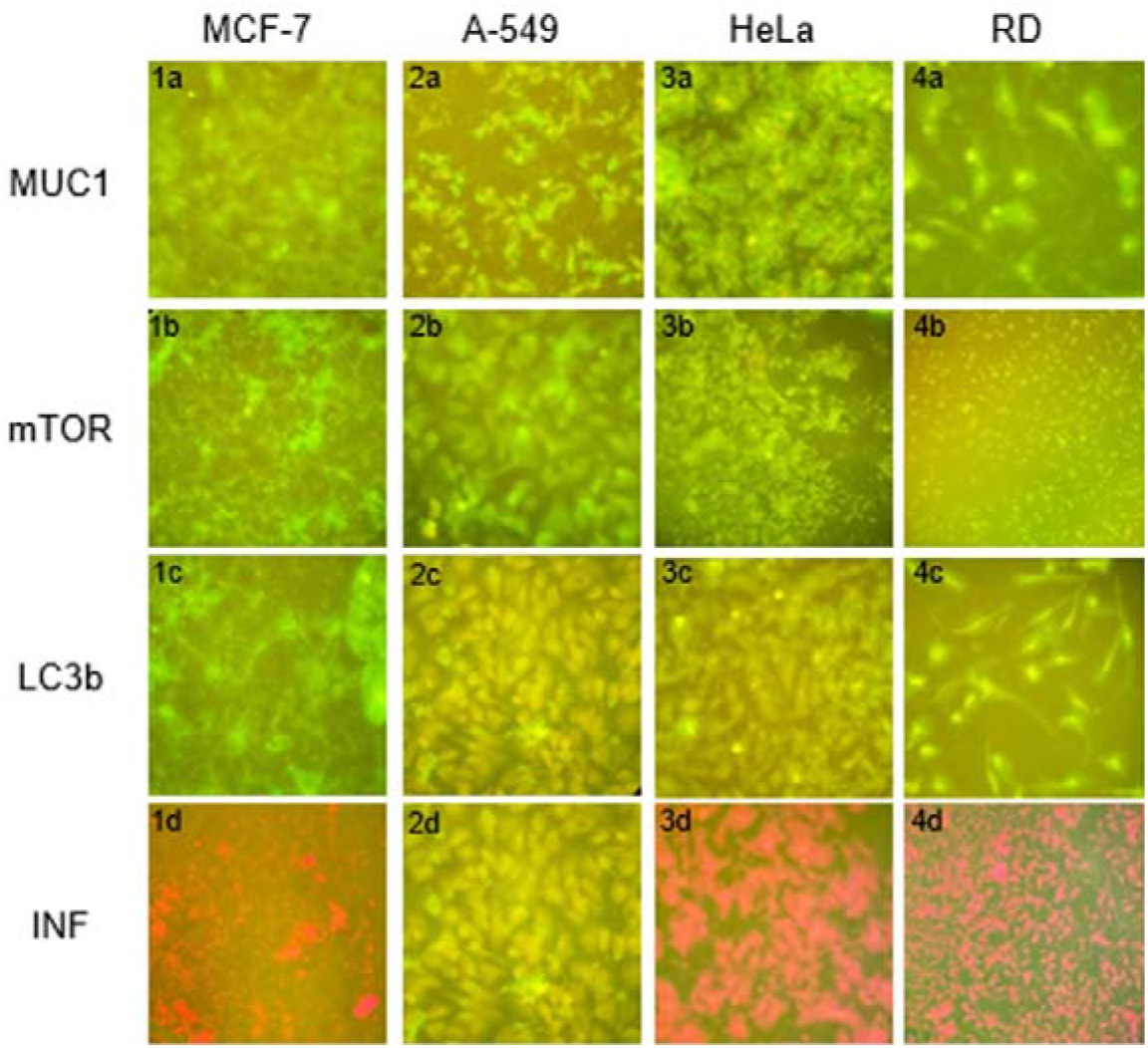
Immunofluorescence staining patterns of influenza negative cells. **1a:** Influenza-negative MCF-7 cells stained with anti-MUC1 antibodies (x40), **2a:** Influenza-negative A-549 cells stained with anti-MUC1 antibodies (x40), **3a:** Influenza-negative HeLa cells stained with anti-MUC1 antibodies (x40), **4a:** Influenza-negative RD cells stained with anti-MUC1 antibodies (x40); **1b:** Influenza-negative MCF-7 cells stained with anti-mTOR antibodies (x40), **2b:** Influenza-negative A-549 cells stained with anti-mTOR antibodies (x40), **3b:** Influenza-negative HeLa cells stained with anti-MUC1 antibodies (x40), **4b:** Influenza-negative RD cells stained with a anti-mTOR antibodies (x10); **1c:** Influenza-negative MCF-7 cells stained with anti-LC3b antibodies (x40), **2c:** Influenza-negative A-549 cells stained with anti-LC3b antibodies (x40), **3c:** Influenza-negative HeLa cells stained with anti-LC3b antibodies (x40), **4c:** Influenza-negative RD cells stained with anti-LC3b antibodies (x40); **1d:** Influenza-negative MCF-7 cells stained with anti-Influenza antibodies (x40), **2d:** Influenza-negative A-549 cells stained with anti-Influenza antibodies (x40), **3d:** Influenza-negative HeLa cells stained with anti-MUC1 antibodies (x40), **4d:**

**Figure 2.**
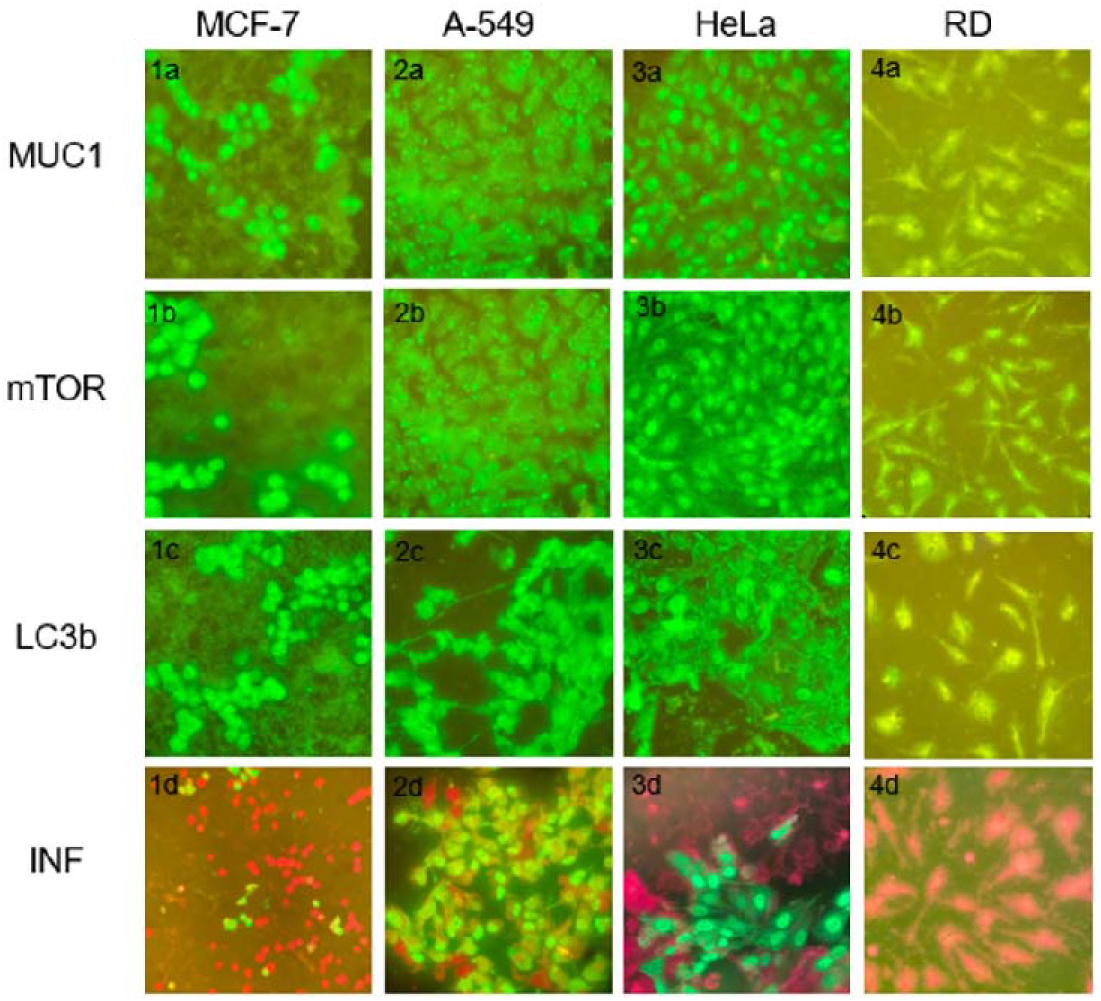
Immunofluorescence staining patterns of influenza positive cells, **1a:** Influenza-positive MCF-7 cells stained with anti-MUC1 antibodies (x40), **2a:** Influenza-positive A-549 cells stained with anti-MUCI antibodies (x40), **3a:** Influenza-positive HeLa cells stained with anti-MUC1 antibodies (x40), **4a:** Influenza-positive RD cells stained with anit-MUC1 antibodies (x40); **lb:** Influenza-positive MCF-7 cells stained with anti-mTOR antibodies (x40), **2b:** Influenza-positive A-549 cells stained with anti-mTOR antibodies (x40), **3b:** Influenza-positive HeLa cells stained with anti-MUC1 antibodies (x40), **4b:** Influenza-positive RD cells stained with a anti-mTOR antibodies (x40) **1c:** Influenza-positive MCF-7 cells stained with anti-LC3b antibodies (x40), **2c:** Influenza-positive A-549 cells stained with anti-LC3b antibodies (x40), **3c:** Influenza-positive HeLa cells stained with anti-LC3b antibodies (x40), **4c:** Influenza-negative RD cells stained with anti-LC3b antibodies (x40); **1d**: Influenza-positive MCF-7 cells stained with anti-Influenza antibodies (x10), **2d:** Influenza-positive A-549 cells stained with anti-Influenza antibodies (x40), **3d:** Influenza-positive HeLa cells stained with anti-MUC1 antibodies (x40), **4d:** Influenza-positive RD cells stained with anti-Influenza antibodies (x40).

In RT-PCR analysis conducted using the supernatants of all cells, the cells inoculated with the virus tested positive for the influenza nucleic acid.

After the isolation of nucleic acid from the cultured cells, gene expressions for *MUC1, MTOR* and *MAP1LC3B* were detected in all inoculated cells except RD cell line. The early expression of all three genes were detected (Ct values 20.01-23.46) in A-549 cells. In the uninfected A-549 cells, gene expressions for all three structures were either undetectable or weak positive (Ct value>30).

The expressions of all three genes in varying degrees were detected in both influenza inoculated or uninoculated MCF-7 and HeLa cell lines. In RD cells inoculated no gene expression regarding the aforementioned three genes were detected, with the inoculated RD cells having expressions of all three genes. All Ct values can be seen in Table 1.

## 4. Discussion

In this study, the role of alternating MUC1 expressions on previously-described autophagy induced by influenza virus has been evaluated in four different cell lines, as MUC1 expressions differ between healthy and cancer cells^7^, and are abundant in some adenocarcinoma cells with being nonexistant or not of importance in mesenchymal cells, as these cells are not organized in a manner to face the lumen in the organism unlike the adenoid cells. The cancers that abundantly express MUC1 are known to have a worse prognosis and higher rate of metastasis^8^.

To support this argument, three adenocarcinoma and one non-adenocarcinoma cell line were used in the study and cell culture experiments were performed both with specific immunofluorescent staining and by measuring gene expirations in cell culture supernatants and isolated cells via RT-PCR. While all immunofluorescent staining results support our argument, the same momentum was not achieved in RT-PCR.

All genes were found to be positive with early Ct values detected only in influenza positive A-549 cells, while gene expressions of *MAP1LC3b, MUC1* and *MTOR* were negative in influenza-negative A-549 cells. However, this result was not obtained in other adenocarcinoma cells. Gene expressions were paradoxically observed in both influenza-positive and influenza-negative MCF-7 and He-La cells. The expressions of genes were negative in influenza-positive RD cells and in influenza-negative RD cells, which was used as a negative control and did not have an adenocarcinoma structure, but is a mesenchymal immature tumor, not known to express MUC1 unlike some other sarcomas^9^, consistent with our results, however whose alveolar histological type is known to express MUC4^10^, which may explain the unexpected influenza-positivity in these cell line, as MUC4 also has sialic acid residues, which could serve as potential influenza receptors^11^.

MUC-1, mTOR and LC3b immunofluorescence stainings were positive in influenza-positive A-549 lung adenocarcinoma cell line, and negative in influenza negative A-549 cells. In addition, fluorescent staining of the virus in these cells is also positive. In nucleic acid tests where quantitative measurement was also performed, influenza virus positivity in A-549 cells was found to be 23.17 Ct. Early entry positivity was detected by nucleic acid test in all MUC-1, mTOR and LC3B in cells inoculated with influenza virus. In addition, all three pathways were found to be negative in influenza virus cells, which were shown to be negative by both PCR and cell culture, both by immunofluorescence tests and nucleic acid tests. Expressions of both MUC1 and autophagy markers mTOR and LC3b were also increased after inoculation with influenza virus in A-549 cell line. The concomitant increase of *MTOR* and *MAP1LC3b* was found to be compatible with the previously described influenza-associated autophagy by Kuss-Duerkop et al^2^. Contrary to our hypothesis, no decrease in *MUC1* gene expressions was detected after inoculation with influenza virus. Although there was no decrease in MUC1 gene expressions in cells, autophagy was realized through an increase in LC3b. Although it is possible to say that the change in gene expressions on DNA corresponds to the transcription level, with the results we have, further studies are required to make a comment on the protein level, as our immunofluorescence results are only reliable in terms of presence of a protein, not providing any information about protein expression levels.

MUC-1, mTOR and LC3B immunofluorescence stainings were found positive in influenza-positive snd negative MCF-7 cells. Influenza virus Ct value was found to be 26.13 in MCF-7 cells, and fluorescent staining of the virus was also positive in these cells. *MUC1, MTOR* and *MAP1LC3B* expressions were found to be positive in influenza-positive and negative MCF-7 cells. Interestingly, in this cell line, *MAP1LC3B, MUC1* and *MTOR* positivity were detected earlier in influenza-negative cells, while gene expressions were decreased in influenza-positive cells. After inoculation with influenza virus, in MCF-7 cells, *MUC1* expressions were decreased, and autophagy markers mTOR and LC3b were also decreased. The concomitant increase of *MTOR* and *MAP1LC3B* has been compatible with influenza-associated autophagy. Consistent with our hypothesis, *MUC1* gene expression was decreased in this cell line after influenza inoculation, but this had no effect on the autophagy process. In this cell line, no relationship was found at the mRNA level in autophagy-related gene expressions.

MUC-1, mTOR and LC3B immunofluorescence stainings were found positive in HeLa cervical adenocarcinoma cell line positive for influenza virus and negative in influenza negative HeLa cells. In addition, fluorescent staining of the virus was found positive in these cells. Influenza virus positivity in He-La cells was found to be 27.92, and fluorescent staining of the virus was also positive in these cells. An increase in Ct values of *MAP1LC3B, MUC1* and *MTOR* and a decrease in the expression of these genes were observed after virus growth in HeLa cell line. These data were not compatible with influenza-associated autophagy in these cell lines. In the evaluation made with cDNA synthesis, *MUC1* and *MTOR* expressions in HeLa cells increased after inoculation and *MAP1LC3B* expressions decreased.

Since the RD cell line is a rhabdomyosarcoma cell of mesenchymal origin, it was used as a negative control alongside other cells. According to the project hypothesis, since this cell is not an adenocarcinoma cell type, no autophagy pathway will be expressed and MUC-1 sialization will not be seen even if influenza growth occurs. As a result of the experiments, it was observed that influenza virus did not grow in immunofluorescent stainings in this cell line, but a positivity was detected in the RT-PCR test, albeit weakly (Ct 33.98). All three pathways were negative in the immunofluorescent staining of influenza inoculated RD cell lines. In nucleic acid tests, the pathways were found to be negative in the cells inoculated with influenza, while the pathways were found to be positive in the influenza negative cells. The opposite of the immunofluorescence test results was obtained with nucleic acid tests.

The most important limitation of this study is the inability to quantitatively determine the protein-level equivalents of changes in gene expressions obtained by qRT-PCR. The comparison of qRT-PCR through 2^-ΔΔCt^ method among groups, as we didn’t obtain the expressions of housekeeping genes, was not possible. Other important limitations are the limited amount of primary antibodies we have and the lack of experience of the study team in the methods used. Our fourth limitation is that the effect of apoptosis^1^, which is known to frequently accompany influenza-related autophagy, on the expression of these genes has not been determined. Despite all this, our study maintains its importance and specificity for the literature in terms of examining MUC1 expressions together with autophagy markers mTOR and LC3b and examining their changes after influenza inoculation.

## 5 Conclusion

As a result, the presence of autophagy pathways activated by influenza virus in adenocarcinoma cells has been demonstrated and is associated with MUC-1 expressions. However, this has not been fully proven by nucleic acid tests. This may be because the nucleic acid tests used in the study only detect gene expression and mRNA formation. In future studies, the expression of these genes at the protein level should be determined through more advanced further testing.

## 8. Acknowledgements

We’d like to hereby sincerely thank to Mr. Sinan Ercan, for his contributions during the conduction of experiments.

